# Sustained Oxygenation Accelerates Diabetic Wound Healing by Simultaneously Promoting Epithelialization and Angiogenesis, and Decreasing Tissue Inflammation

**DOI:** 10.1101/2021.04.06.438689

**Authors:** Ya Guan, Hong Niu, Zhongting Liu, Yu Dang, Jie Shen, Mohamed Zayed, Liang Ma, Jianjun Guan

## Abstract

Non-healing diabetic wound is one of the most common complications for diabetic patients. Chronic hypoxia is among the prominent factors that delay the wound healing process. Therefore, sustained oxygenation to alleviate hypoxia is hypothesized to promote diabetic wound healing. Yet it cannot be achieved by current clinical approaches including hyperbaric oxygen therapy. Herein, we developed a sustained oxygenation system consisting of oxygen-release microspheres and a reactive oxygen species (ROS)-scavenging hydrogel. The hydrogel was used to capture the ROS that is elevated in the diabetic wounds, and that may be generated due to oxygen release. The sustainedly released oxygen augmented survival and migration of keratinocytes and dermal fibroblasts; promoted angiogenic growth factor expression, and angiogenesis in the diabetic wounds; and decreased M1 macrophage density. These effects led to a significant increase of wound closure rate. These findings reveal that sustained oxygenation alone without using drugs is capable of healing diabetic wounds.

## Introduction

Diabetes is a chronic metabolic disorder affecting 34.2 million people across the United States (*1*). One of the common complications of diabetes is diabetic foot ulcers (DFU). Roughly 25% of diabetic patients experience DFU (*2*), which causes pain, lowered vitality and possible need for limb amputation. Despite advances in conservative medical care such as wound off-loading, wound debridement, and infection control (*3*), these approaches have inconsistent outcomes and are associated with undesired side effects (*3-5*). Therefore, stable and safe treatments for chronic, non-healing wounds need to be developed.

Cutaneous diabetic wound healing is a complex process with three overlapping phases: inflammation, proliferation, and remodeling (*6-8*). Immediately after the injury, impaired vasculature impedes oxygen delivery to the wound, creating a hypoxic environment around the wound (*6, 9, 10*). This hypoxia is exacerbated by the recruitment of inflammatory cells with high oxygen consumption (*11, 12*). Though acute hypoxia promotes cell proliferation and initiates tissue repair, long-term oxygen deprivation in chronic wounds impairs the healing process by inhibiting angiogenesis, re-epithelialization, and extracellular matrix (ECM) synthesis (*13*). Thus, enhanced wound tissue oxygenation is key to chronic wound healing.

In the past few decades, diabetic wound healing has been clinically facilitated by oxygenation, particularly hyperbaric oxygen therapy (HBOT) (*14, 15*). HBOT delivers 100% oxygen at 2-3 atmospheres for 1-2 hours per treatment to patients with DFU. In some cases, HBOT promoted healing of diabetic wounds after 40 or more treatments (*16, 17*), while it did not show obvious effect in other studies (*14, 18*). Overall, the therapeutic efficacy of HBOT is widely considered to be inconsistent and unsatisfactory (*19, 20*). This has mainly resulted from the incapability of HBOT to continuously provide sufficient oxygen to the wounds as the oxygen content in the poorly vascularized wounds decreases quickly following the treatment (*21*). Moreover, as a systemic oxygen delivery strategy, HBOT may lead to risks of tissue hyperoxia such as oxygen toxicity seizure (*21, 22*).

To address the limitations of systemic oxygenation, several previous reports have developed oxygen-generating systems that are implanted locally to increase the oxygen concentration in wound beds. These oxygen-generating systems were based on H_2_O_2_ (*23*), calcium peroxide (*24, 25*) and perfluorocarbon (*26*). The localized oxygenation can avoid systemic hyperoxia while accelerating chronic wound healing. However, these oxygenation systems typically release oxygen for only 3-6 days (*24, 26*), which is not long enough for diabetic wound healing. In addition, the systems cannot quickly release sufficient oxygen to relieve hypoxia (*24, 25*). Oxygen is required for crucial events such as angiogenesis, granulation, re-epithelialization, and ECM synthesis that often take two weeks or longer (*9, 27, 28*). Hence, it is critical to develop oxygen-generating systems to continuously oxygenate the wound bed to accelerate healing.

Herein, we designed an oxygen-generating system based on oxygen-release microspheres (ORMs) and an injectable, fast gelation, and ROS-scavenging hydrogel (ROSS gel). The microspheres were able to quickly release enough oxygen to support cell survival under hypoxia, and sustainedly release oxygen for at least 2 weeks. Unlike most current oxygen-generating systems that release toxic H_2_O_2_ into tissue environment first, and rely on the H_2_O_2_ decomposition to release oxygen, the designed oxygen-release microspheres directly release oxygen. The injectable and fast gelation hydrogel was used as carrier to deliver the microspheres into wounds, and quickly immobilize them in the wounds with high tissue retention rate. The hydrogel had relatively high water content, enabling it to maintain a moisture environment surrounding the wound (*29, 30*). The ROS-scavenging property allows the oxygen-generating system to capture excessive ROS in the diabetic wounds. While ROS plays a crucial role in regulating biological and physiological processes, elevated ROS in diabetic wounds damages cells (*31*). In this study, we assessed the therapeutic efficacy of the oxygen-generating system in promoting wound closure. We also elucidated the underlying mechanisms of how sustained released oxygen promotes diabetic wound healing. While previous studies have shown that short-term oxygen release facilitated wound closure, the underlying mechanisms are not fully clear, and are mainly attributed to wound angiogenesis (*32*). We comprehensively evaluated the effect of sustainedly released oxygen on skin cell survival, migration, and paracrine effects; intracellular oxygen content; prosurvival pathways; tissue angiogenesis; and tissue inflammation and oxidative stress.

## Results

### Microspheres capable of sustainedly releasing molecular oxygen

The ORMs were designed to have core-shell structure (**Fig. 1A**). The core was a stable polyvinylpyrrolidone (PVP)/hydrogen peroxide (H_2_O_2_) complex. The shell was a bioeliminable and bioconjugatable poly(N-isopropylacrylamide-co-2-hydroxyethyl methacrylate-co-acrylate-oligolactide-co-N-acryloxysuccinimide) (abbreviated as poly(NIPAAm-co-HEMA-co-AOLA-co-NAS)) (**Fig. S1**). The microspheres had a diameter of ∼5 µm (**Fig. 1B**). The core-shell structure was confirmed by fluorescent images (**Fig. 1C**). The shell of the microspheres was conjugated with catalase in order to timely convert the H_2_O_2_ in released PVP/H_2_O_2_ into molecular oxygen, and avoid H_2_O_2_ to exit the microspheres causing toxicity concerns. A layer of catalase was conjugated on the shell as evidenced by the fluorescence signal of FITC-labeled catalase (**Fig. 1D**).

**Fig. 1.**
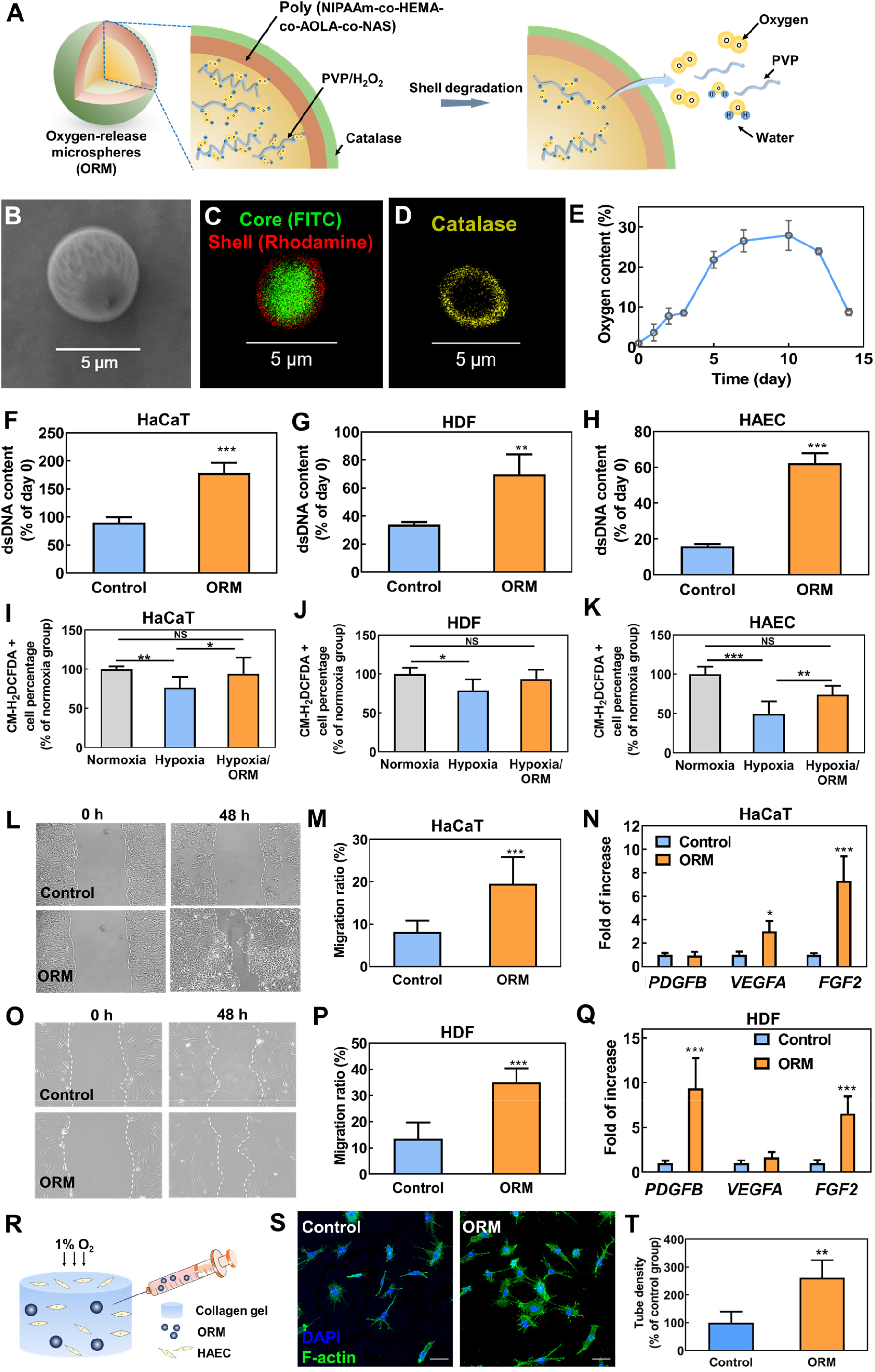
Oxygen-release microspheres provide continuous oxygenation and regulate cell behaviors in vitro. **(A)** Schematic illustration of the oxygen-release microsphere and its oxygen-release mechanism. **(B)** SEM images of the microspheres. (**C**) Fluorescent images of the microspheres. The core and shell of the microspheres were labeled with FITC (core) and rhodamine (shell), respectively. (**D**) Fluorescent images of the microspheres after catalase conjugation. Catalase was pre-labeled with FITC. (**E**) Oxygen release kinetics of the gel/ORM construct for 14 days. The oxygen content was determined by measuring the fluorescence intensity of an O_2_-sensitive fluorophore Ru(Ph_2_phen_3_)Cl_2_ and converting into oxygen level using a calibration curve. n = 8. (**F**) dsDNA content of keratinocytes (HaCaT cells) cultured under hypoxic condition for 5 days. n = 5. (**G**) dsDNA content of dermal fibroblasts (HDF) cultured under hypoxic condition for 5 days. n = 5. (**H**) dsDNA content of endothelial cells (HAEC) cultured under hypoxia condition for 7 days. n = 3; (**I**) ROS content in keratinocytes (HaCaT cells) cultured under normoxia, hypoxia, and hypoxia with microspheres for 3 days. n = 10. (**J**) ROS content in dermal fibroblasts (HDF) cultured under normoxia, hypoxia, and hypoxia with microspheres for 5 days. n = 6. (**K**) ROS content in endothelial cells (HAEC) cultured under normoxia, hypoxia, and hypoxia with microspheres for 3 days. n = 8. (**L, M**) Migration of keratinocytes cultured under hypoxia for 48 hours and corresponding quantifications. n = 4. (**N**) Gene expression of *PDGFB, VEGFA* and *FGF2* in keratinocytes cultured under hypoxic conditions for 48 hours. n ≥ 3. (**O, P**) Migration of dermal fibroblasts cultured under hypoxia for 48 hours and corresponding quantifications. n = 4. (**Q**) Gene expression of *PDGFB, VEGFA* and *FGF2* in dermal fibroblasts cultured under hypoxic conditions for 48 hours. n ≥ 3. (**R**) Schematic illustration of the in vitro tube formation assay; (**S**) Endothelial cells tube formation. Endothelial cells were cultured under hypoxia for 16 hours and stained with F-actin (green) and DAPI (blue). Scale bar = 50 µm. (**T**) Quantification of tube density. *p<0.05, **p<0.01, ***p<0.001.

To determine oxygen release kinetics, we used an oxygen-sensitive luminophore Ru(Ph_2_phen_3_)Cl_2_ whose fluorescence intensity is linearly proportional to the oxygen content (*34, 35*). The ORMs were able to continuously release oxygen during the 2-week experimental period (**Fig. 1E**). The oxygen level reached above 5% after 2 days of release. It was maintained above 10% from day 3 to day 14. After 2-week study period, the release medium contained less than 10 µM H_2_O_2_ as measured by a quantitative peroxide assay kit. This concentration will not cause cell apoptosis (*36, 37*).

### Continuous oxygenation of skin cells by released oxygen to promote skin cell survival, migration and paracrine effects, and endothelial lumen formation under hypoxia in vitro

In diabetic wounds, hypoxia is one of the major factors that compromise cell survival and migration, leading to slow wound healing (*9, 35, 38*). To evaluate whether oxygen released from ORMs was able to improve cell survival under hypoxia, we incubated human keratinocytes (HaCaT cells), human dermal fibroblasts (HDFs), and human arterial endothelial cells (HAECs) respectively with the ORMs under 1% oxygen condition. All 3 cell types exhibited a significantly higher double-stranded DNA (dsDNA, for live cells) content than the corresponding control groups without ORMs (**Fig. 1, F to H**, p<0.001 for HaCaT cells and HAEC, p<0.01 for HDF). These results demonstrate that the released oxygen effectively increased skin cell survival under hypoxia. One of the concerns of oxygen treatment is overproduction of ROS. To characterize ROS expression in the 3 cell types, we used an ROS-sensitive dye CM-H_2_DCFDA to stain the cells cultured under normoxia, hypoxia (1% oxygen condition), and hypoxia with addition of ORMs (hypoxia/ORM group). The released oxygen in the hypoxia/ORM group significantly increased ROS level in the 3 cell types compared to the hypoxia group (**Fig. 1, I to K, Fig. S2-4**). However, the ROS level was similar to the normoxia group (p>0.05 for all cell types). These results show that the released oxygen did not overproduce ROS leading to cell apoptosis.

To determine whether the released oxygen can increase the migration of keratinocytes and dermal fibroblasts, a scratch assay was performed. After 48 hours of incubation under 1% oxygen condition, the migration rates of HaCaT cells and HDFs were significantly higher in the ORM groups than the control groups without ORMs (**Fig. 1, L, M, O and P**, p<0.001). These results demonstrate that the released oxygen promoted skin cell migration under hypoxia.

We further elucidated the effect of released oxygen on skin cell expression of angiogenic growth factors under hypoxic condition. For HaCaT cells, the treatment with ORMs significantly increased the expression of vascular endothelial growth factor-A (*VEGFA*, p<0.05) and basic fibroblast growth factor (*FGF2*, p<0.001) (**Fig. 1N**). For HDFs, the released oxygen significantly augmented the expression of platelet-derived growth factor-B (*PDGFB*, p<0.001) and *FGF2* (p<0.001) (**Fig. 1Q**). These growth factors play critical roles in re-epithelialization and angiogenesis during cutaneous wound healing (*39-42*).

In chronic wounds, the hypoxic environment impairs angiogenesis, resulting in delayed wound healing (*43*). To evaluate the potential of released oxygen in promoting angiogenesis under hypoxia, in vitro endothelial tube formation assay was conducted (**Fig. 1R**). The ORM was injected into the 3D collagen constructs seeded with HAECs. Following 16 h of culture under 1% oxygen condition, the HAECs assembled significantly greater number of lumens, with ∼2.5 fold of increase in lumen density compared with the control group (**Fig. 1, S and T**, p<0.01).

### Continuous oxygenation by released oxygen to elevate intracellular oxygen content and ATP content, and activate Erk1/2 and HO-1 signaling

To explore the underlying mechanisms that the continuous oxygenation by released oxygen increased skin cell survival and migration, we measured the intracellular oxygen content in HaCaT cells using electron paramagnetic resonance (EPR) (**Fig. 2A**). After 24 hours of treatment with ORMs under 1% oxygen condition, the intracellular oxygen content was 2.5 times of the cells without ORM treatment (**Fig. 2, B and C**). To determine whether the elevated intracellular oxygen content promoted cellular energy production, intracellular adenosine triphosphate (ATP) level in HaCaT cells was measured. The ATP level in the group treated with ORMs was significantly increased compared to that in the group without ORM treatment (**Fig. 2D**, p<0.01). These results demonstrate that the increased cell survival and migration by continuous oxygenation is associated with increased intracellular oxygen content and energy regeneration.

**Fig. 2.**
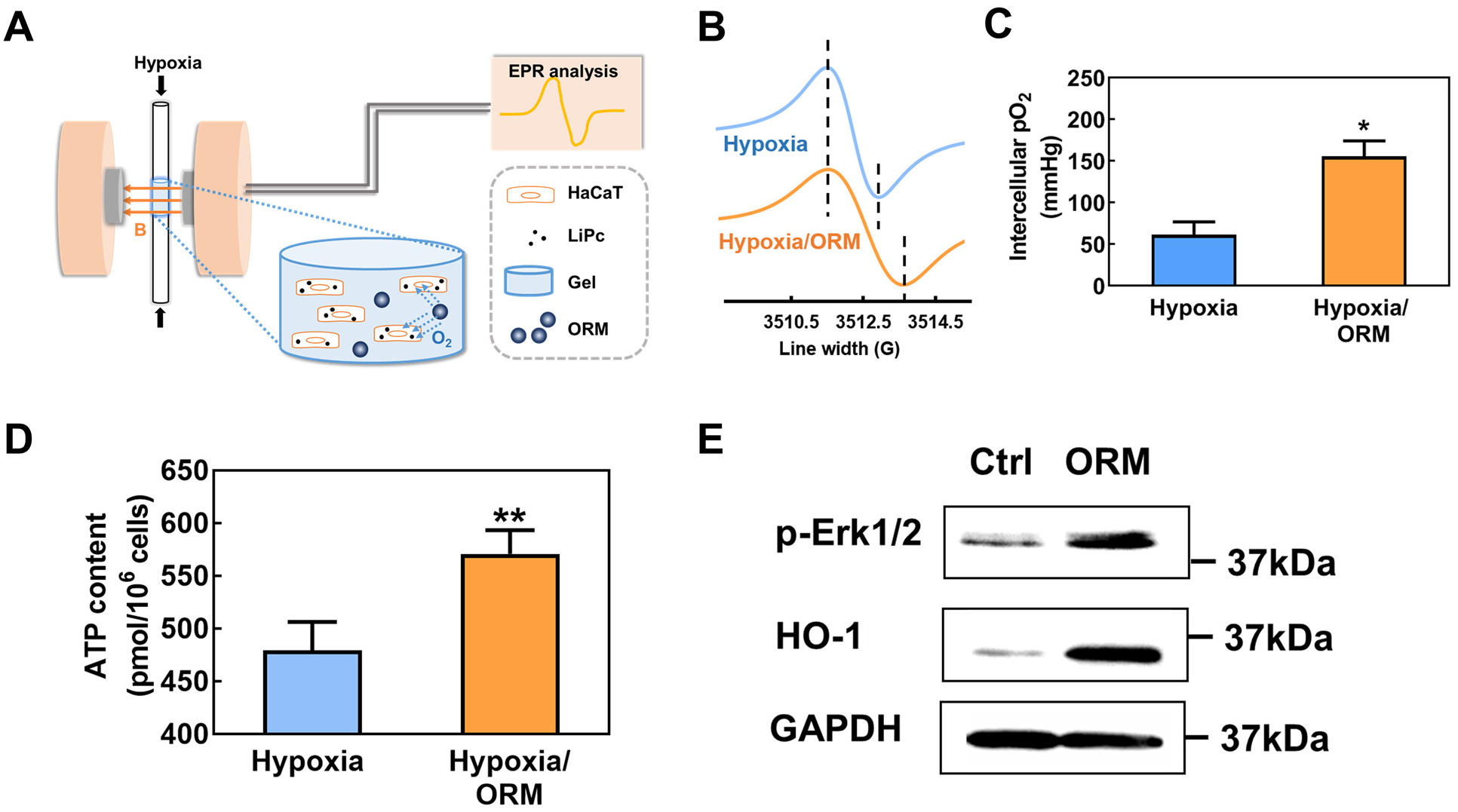
Oxygen-release microspheres elevate intracellular oxygen content, ATP content and activate HO-1 and Erk 1/2 pathways under hypoxia. (**A-C**) Intracellular oxygen content measured by electron paramagnetic resonance (EPR). Keratinocytes (HaCaT cells) were incubated with lithium phthalocyanine nanoparticles for endocytosis, and then cultured under 1% oxygen conditions (with or without ORM) for 24 hours. n = 3 for each group. *p<0.05. (**D**) Intracellular ATP content in HaCaT cells measured by an ATP assay kit. Cells were cultured under 1% oxygen conditions (with or without ORM) for 24 hours. **p<0.01 (**E**) Immunoblotting of HO-1 and phosphorylated Erk 1/2 in dermal fibroblasts cultured under hypoxic conditions for 48 hours. GAPDH serves as a loading control.

The mitogen-activated protein kinases (MAPK) signaling pathway is responsible for regulating cell survival and migration (*44*). We performed western blot analysis using HDF to determine whether extracellular signal-regulated kinases (Erk1/2) axis, one of the major cascades of MAPK pathway, was activated in response to the released oxygen. The results demonstrated that Erk1/2 phosphorylation was increased after the cells were treated with ORMs under 1% oxygen condition (**Fig. 2E**). In addition, we investigated whether heme oxygenase-1 (HO-1), a stress-inducible and cytoprotective protein, was upregulated after treatment with released oxygen under hypoxia. In the ORM group, the HO-1 expression was more pronounced than the control group (**Fig. 2E**). The above results reveal that continuous oxygenation by released oxygen activated Erk1/2 and HO-1 pathway leading to enhanced cell survival and migration.

### Injectable, thermosenstive, and ROS-scavenging hydrogel to deliver oxygen-release microspheres and protect skin cells under oxidative stress

To deliver the ORMs to diabetic wounds and largely retain them in the tissue, we synthesized an injectable and thermosensitive hydrogel with fast gelation rate. The hydrogel was also designed to have ROS-scavenging capability so as to capture upregulated ROS in the diabetic wounds to protect skin cells from ROS-induced apoptosis. This hydrogel may also eliminate ROS that may be generated due to excessive oxygen release. The hydrogel was synthesized by copolymerization of NIPAAm, HEMA and 4-(acryloyloxymethyl)-phenylboronic acid pinacol ester (**Fig. 3, A and B, Fig. S5**). The hydrogel solution (6 wt%) had a gelation temperature of 17°C, and was injectable at 4°C. After being transferred into a 37°C water bath, the hydrogel solution solidified within 6 seconds to form a solid hydrogel (**Fig. 3C**). The mixture of hydrogel solution and ORMs (40 mg/mL) remained injectable at 4°C, and fast gelling (6 seconds) at 37°C (**Fig. 3C**).

**Fig. 3.**
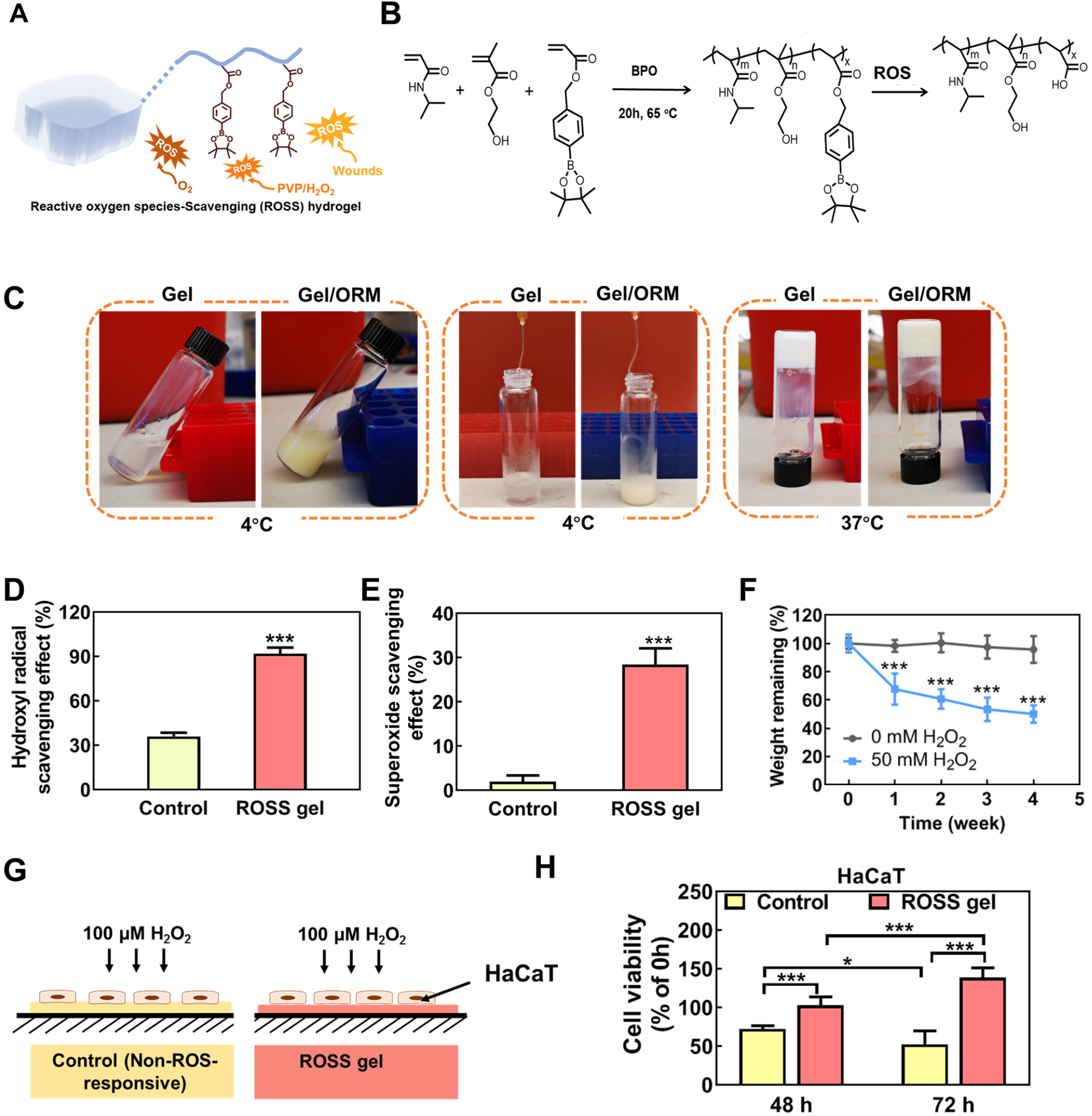
Reactive oxygen species-scavenging hydrogel protects skin cells under oxidative stress by consuming hydroxyl radicals and superoxides. (**A**) ROSS hydrogel capable of scavenging reactive oxygen species in chronic wound tissues or generated from released oxygen or PVP/H_2_O_2_. (**B**) Synthesis and ROS-scavenging mechanism of ROSS hydrogel. (**C**) Injectability and gelation of ROSS hydrogel (Gel) and Gel/ORM construct. (**D**) Scavenging effect on hydroxyl radicals of ROSS gel and non-ROS responsive gel (n=4). (**E**) Scavenging effect on superoxide generation determined by pyrogallol assay (n=3). (**F**) Degradation of ROSS gel in DPBS at 37 °C for 4 weeks with 0 and 50 mM H_2_O_2_. (**G**) Schematic illustration of an in vitro model for skin cell survival on gels under 100 μM H_2_O_2_ to mimic the in vivo cell environment under oxidative stress. Non-ROS-responsive gel was used as a control. (**H**) Cell viability of HaCaT at 48 hours and 72 hours on control gel and ROSS gel normalized to the initial viability on each gel at 0 hour. n≥ 6, *p<0.05, ***p<0.001.

The ROS-scavenging capability of the hydrogel was evaluated in terms of consumption of hydroxyl radical (HO•) and superoxide (O_2_•^-^), and hydrogel weight loss induced by H_2_O_2_. After incubating with HO• for 1 day, the hydrogel scavenged remarkably higher HO• than the non-ROS-responsive, control hydrogel (**Fig. 3D**, p<0.001). Similarly, the hydrogel eliminated 8-fold more O_2_•^-^ than the control hydrogel after incubating for 1 day (**Fig. 3E**, p<0.001). The hydrogel gradually lost weight in H_2_O_2_ solution during the 4-week experimental period (**Fig. 3F**). In contrast, the hydrogel did not show substantial weight loss in Dulbecco’s phosphate-buffered saline (DPBS) without H_2_O_2_.

To evaluate efficacy of the hydrogel in protecting skin cell survival under pathological oxidative stress, HaCaT cells were seeded on the hydrogel surface in the presence of 100 μM H_2_O_2_, and the cell viability was quantified (**Fig. 3G**). The non-ROS-responsive hydrogel was used as control. We found that HaCaT cells seeded on the hydrogel did not undergo substantial apoptosis in 100 μM H_2_O_2_ at 48 hours, and even proliferated at 72 hours. However, HaCaT cells on the non-ROS-responsive hydrogel had significantly lower viability than the ROS-responsive hydrogel (p<0.001 at 48 hours and 72 hours), with only ∼50% viability at 72 hours (**Fig. 3H**). These results demonstrate that the ROS-responsive hydrogel was able to protect keratinocytes from apoptosis caused by pathological oxidative stress.

### Accelerated diabetic wound closure by continuous oxygenation and ROS scavenging

To determine whether continuous oxygenation and ROS scavenging can accelerate diabetic wound healing, we administered the hydrogel with ORMs (Gel/ORM group) onto full thickness excisional wounds on diabetic mice (**Fig. 4A**). Wounds without treatment (No-treatment group) and wounds treated with hydrogel alone (Gel group) were used as controls. The wounds treated with Gel/ORM had a greater closure rate than the No-treatment and Gel groups (**Fig. 4B**). By day 16, the wound size in the Gel/ORM group was reduced to 10.7%, significantly smaller than that in the Gel (30.4%) and No-treatment groups (52.2%) (p<0.01, **Fig. 4C**). Notably, the wound size in the Gel group was significantly smaller than the No-treatment group, demonstrating that the Gel alone promoted wound closure.

**Fig. 4.**
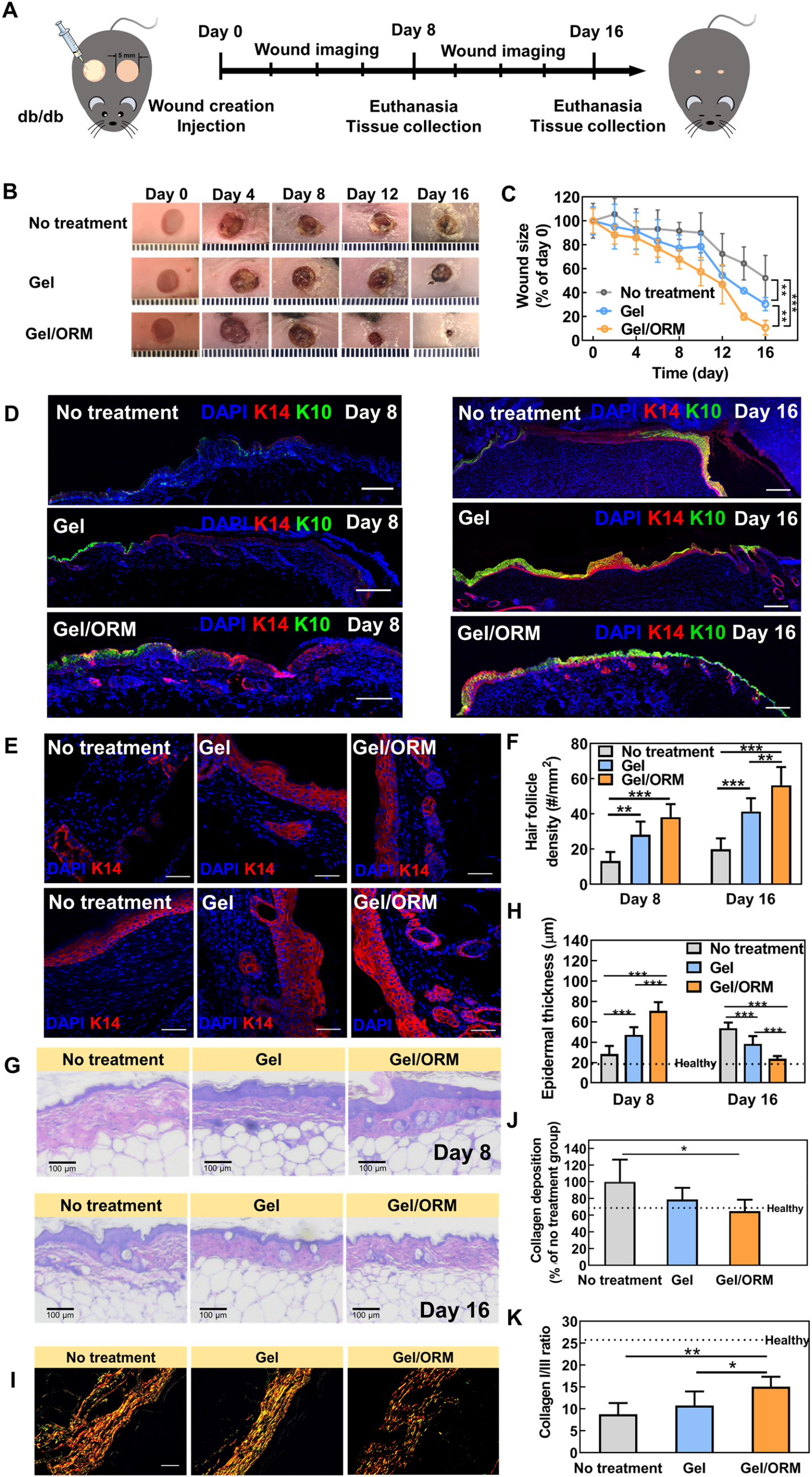
Oxygen-release microspheres encapsulated in ROSS hydrogel accelerate wound healing in db/db mice. (**A**) Schematic illustration of the design of animal experiments to test the therapeutic effect of gel/ORM in a db/db mouse model. (**B**) Representative images of the wounds treated with or without gel and microspheres for 16 days. (**C**) Wound closure rate for 16 days post wounding. Wound size at each time point was normalized to day 0. n ≥ 8. (**D**) Immunohistochemical staining of cytokeratin 10 (K10, green) and cytokeratin 14 (K14, red) of the wounds at day 8 and day 16. Scale bar = 200 µm. (**E**) Immunohistochemical staining of K14 (red) in the wounded region at 8 and 16 days postwounding. Scale bar = 50 µm. (**F**) Quantification of hair follicle density at day 8 and day 16. (**G**) H&E staining of the wounded skin at day 8 and day 16. (**H**) Quantification of epidermal thickness of the wounded skin. (**I**)Picrosirius red staining of the wounded skin at day 16. Scale bar = 50 µm. (**J**) Quantification of total collagen deposition at day 16. (**K**) Quantification of collagen I/III ratio at day 16. *p<0.05, **p<0.01, ***p<0.001.

Re-epithelialization is a crucial process in wound healing. To determine the re-epithelialization process during the treatment, staining of cytokeratin 14 (K14) and cytokeratin 10 (K10) was performed on wounds harvested at days 8 and 16. The two markers are for basal keratinocytes and spinous keratinocytes, respectively. At day 8, the wound gap was nearly enclosed with a complete layer of basal keratinocytes (K14+) in the Gel/ORM group, while keratinocyte migration was much slower in the No-treatment and Gel groups (**Fig. 4D**). In addition, the Gel/ORM group formed a more matured spinous layer composed of K10+ keratinocytes compared to the No-treatment and Gel groups. At day 16, the migration of basal keratinocytes was complete, and the proliferation and differentiation were ongoing in the No-treatment and Gel groups. In contrast, the re-epithelialization was fully completed in the Gel/ORM group, as evidenced by a clear stratified epithelium (**Fig. 4D**). K14 staining was performed for the hair follicles in the basal layer of epidermis (**Fig. 4E**). A significant increase in hair follicle density was found for the Gel/ORM group than No-treatment and Gel groups at both day 8 and day 16 (**Fig. 4F**, p<0.01). These results demonstrate that the continuous oxygenation by released oxygen, and ROS scavenging effectively accelerated keratinocyte migration and hair follicle formation, consequently promoted re-epithelialization of the diabetic wounds.

Epidermal thickness is another indicator of the healing process. It increases during the initial inflammatory and proliferative stages of the wound healing process, then decreases during the remodeling stage (*45*). At day 8, the wounds treated with Gel/ORM exhibited the thickest epidermis. It became the thinnest at day 16 (**Fig. 4, G and H**, p<0.001), suggesting the continuous oxygenation and ROS scavenging facilitated the wound healing process from inflammatory and proliferative stages to the remodeling stage.

To determine whether the accelerated wound closure was associated with scar formation, picrosirius red staining was performed for collagens in the wounds. The Gel/ORM group had a significantly lower total collagen content than the No-treatment group at day 16 (**Fig. 4, I and J**, p<0.05). In addition, a significantly higher collagen I/III ratio was found for the Gel/ORM group than the No-treatment and Gel groups (**Fig. 4K**, p<0.05). This collagen I/III ratio is more close to that of the skin without injury than the control groups. The reduced collagen deposition and higher collagen I/III ratio demonstrate that the enhanced wound healing did not induce scar formation (*46, 47*).

### Promoted cell proliferation and metabolism by continuous oxygenation and ROS scavenging in diabetic wounds

To understand the role of continuous oxygenation and ROS scavenging in diabetic wound healing at cellular level, cell proliferation and metabolism in the wounds were examined. The treatment with ROS-scavenging hydrogel significantly increased the density of Ki67+ proliferating cells compared to the No-treatment group at both day 8 and day 16. The administration of Gel/ORM further significantly increased the density of proliferating cells with more pronounced increase at day 16 (**Fig. 5, A and C**). Continuous oxygenation and ROS scavenging also increased skin cell metabolic rate as judged by PGC1α+ cell density (**Fig. 5, B and D**) (*48*). Compared with No-treatment group, the Gel group exhibited significantly higher density of PGC1α+ cells. Release of oxygen in the Gel/ORM group more greatly increased the PGC1α+ cell density at both time points.

**Fig. 5.**
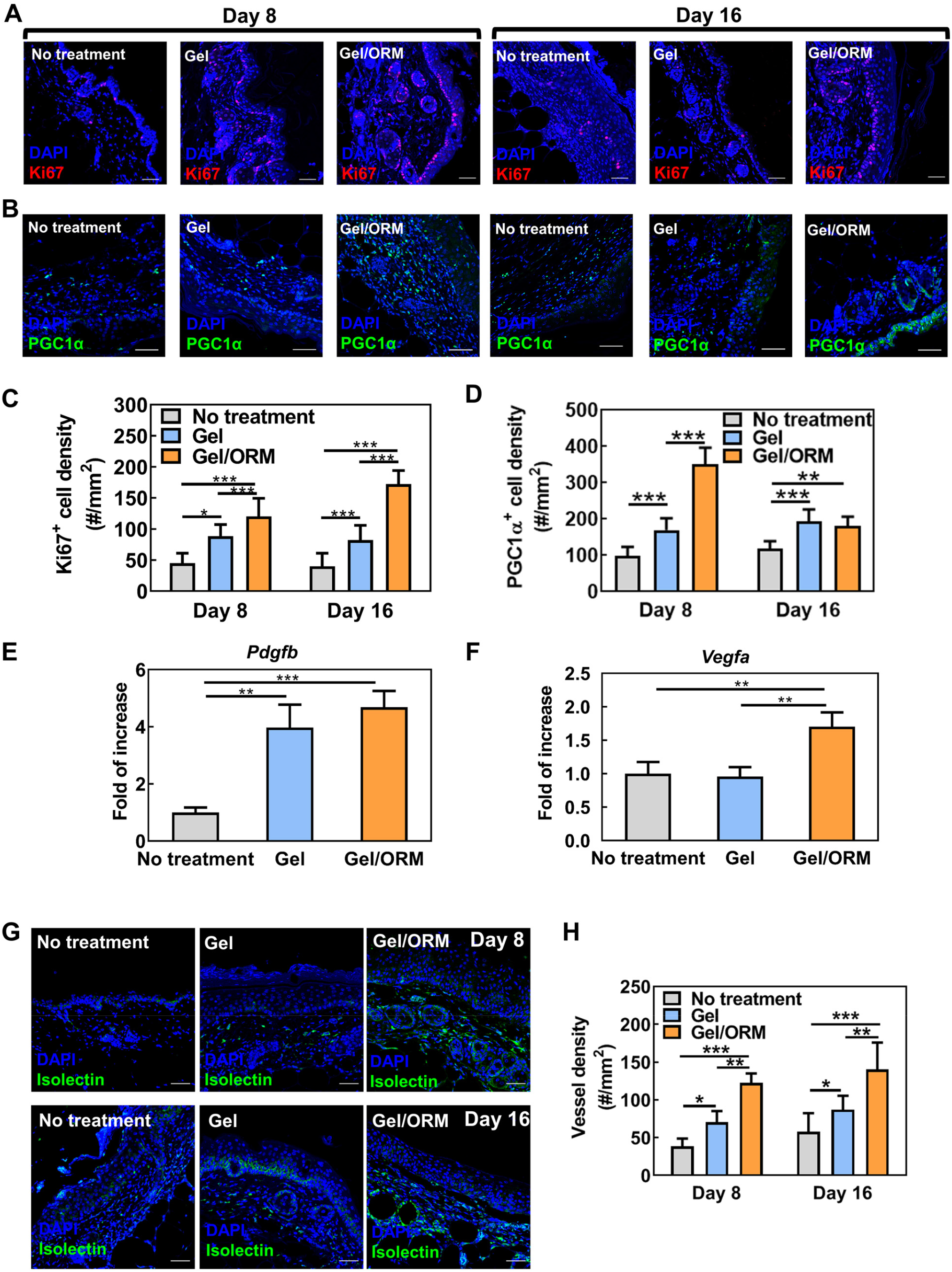
Continuous oxygenation and ROS scavenging promoted cell proliferation and metabolism, and stimulated angiogenic growth factor expression and angiogenesis in diabetic wounds. (**A**) Immunohistochemical staining of Ki67 (red) in the wounded region at 8 and 16 days postwounding. Scale bar = 50 µm. (**B**) Immunohistochemical staining of PGC1-α (green) in the wounded region at 8 and 16 days postwounding. Scale bar = 50 µm. (**C**) Quantification of Ki67 positive cell density. (**D**) Quantification of PGC1-α positive cell density. (**E, F**) Gene expression of *Pdgfb* (**E**) and *Vegfa* (**F**) from the tissue lysates extracted from the wounded skin at day 8. n ≥ 4. (**G**) Immunohistochemical staining of isolectin (green) in the wounded region at 8 and 16 days postwounding. Scale bar = 50 µm. Nuclei were stained with DAPI in all immunohistochemical staining images. (**H**) Quantification of vessel density. *p<0.05, **p<0.01, ***p<0.001.

### Stimulated angiogenic growth factor expression, and angiogenesis by continuous oxygenation and ROS scavenging in diabetic wounds

To determine whether continuous oxygenation and ROS scavenging affects the expression of angiogenic growth factors in the diabetic wounds, real-time RT-PCR was performed using the wound tissues 8 days after surgery (**Fig. 5, E and F**). Among different angiogenic growth factors, the *Pdgfb* expression was significantly increased in the Gel and Gel/ORM groups (**Fig. 5 E**, p<0.01). The Gel/ORM group had substantially greater expression than the Gel group (p>0.05). Notably, the Gel/ORM group exhibited significantly higher *Vegfa* expression than the Gel and No-treatment groups (**Fig. 5F**, p<0.01).

To evaluate whether continuous oxygenation and ROS scavenging stimulated angiogenesis in the diabetic wounds, the capillary densities in the Gel/ORM, Gel and No-treatment groups were quantified at middle (day 8) and late (day 16) stages of the wound healing. At both stages, the Gel group had significantly higher density of capillaries than the No-treatment group, demonstrating that ROS-scavenging promoted diabetic wound angiogenesis (**Fig. 5, G and H**). The simultaneous ROS-scavenging and continuous oxygenation in the Gel/ORM group further significantly stimulated angiogenesis (**Fig. 5, G and H**).

### Alleviated oxidative stress, inflammation, and pro-inflammatory cytokine expression by continuous oxygenation and ROS scavenging in diabetic wounds

The use of ROS-scavenging hydrogel decreased ROS content in the diabetic wounds (**Fig. 6, A and C**). At both day 8 and 16, the ROS+ cell density was remarkably lower in the Gel group than the No-treatment group (p<0.001). Continuous oxygenation of the diabetic wounds may lead to the formation of ROS although the ROS scavenging hydrogel has the capability to capture it. At day 8 and 16, the Gel and Gel/ORM groups showed similar density of ROS+ cells (**Fig. 6, A and C**, p>0.05), demonstrating that the hydrogel can efficiently scavenge ROS even if the continuous oxygenation induced ROS formation.

**Fig. 6.**
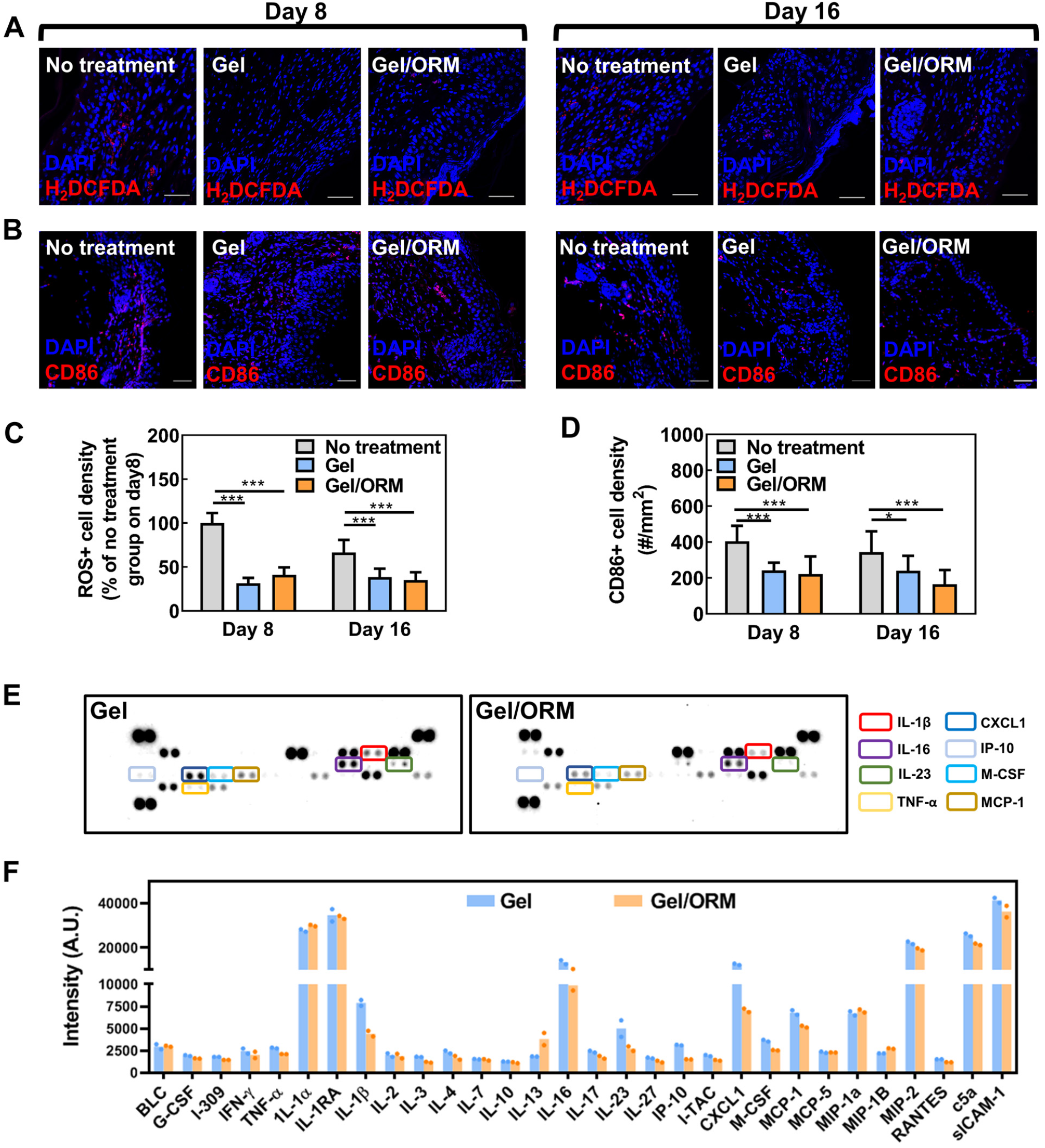
Continuous oxygenation and ROS scavenging alleviated oxidative stress, inflammation, and pro-inflammatory cytokine expression in diabetic wounds. (**A**) Immunohistochemical staining of CM-H_2_DCFDA (red) at the wounded site at day 8 and day 16. Scale bar = 50 µm. (**B**) Immunohistochemical staining of CD86 (red) at the wounded site at day 8 and day 16. Scale bar = 50 µm. Nuclei were stained with DAPI in all immunohistochemical staining images. (**C**) Quantification of CD86 positive cell density. n ≥ 8. (**D**) Quantification of ROS positive cell density. The results were normalized to the ROS+ cell density in no-treatment group on day 8. *p<0.05, ***p<0.001. (**E**) Cytokine array analysis of the pro-inflammatory cytokine level in the wound tissues 8 days after treatment. (**F**) Quantitative summary of cytokine array analysis in (**E**).

To elucidate the impact of continuous oxygenation and ROS scavenging on the inflammatory response in diabetic wound, pro-inflammatory M1 macrophages in the tissue were characterized. The M1 macrophage density (CD86+) in the Gel and Gel/ORM groups were significantly lower than the No-treatment group (**Fig. 6, B and D**, p<0.001 at day 8, and p<0.05 at day 16). Notably, the Gel and Gel/ORM groups had similar M1 macrophage cell density. These results demonstrate that ROS scavenging by the hydrogel decreased diabetic wound inflammation.

To further reveal the role of released oxygen in tissue inflammation, pro-inflammatory cytokine expressions in the diabetic wounds were determined (**Fig. 6, E and F**). Compared with Gel group, Gel/ORMs groups reduced the expression of various pro-inflammatory cytokines, such as interleukin 1 beta (IL-1β), tumor necrosis factor (TNF-α), interferon gamma (IFN-γ) and chemokine ligand 1(CXCL1). These results suggest that continuous oxygenation has potential to decrease inflammation during diabetic wound healing.

### Upregulated pErk1/2 and HO-1 expressions by continuous oxygenation in diabetic wounds

As we have shown that the oxygenation of keratinocytes by released oxygen upregulated the p-Erk1/2 and HO-1 expressions in vitro, we evaluated whether continuous oxygenation upregulated these expressions in the diabetic wounds. Consistent with the in vitro findings, the p-Erk1/2 and HO-1 expressions were substantially upregulated in the Gel/ORM group at day 8 and 16 (**Fig. 7**).

**Fig. 7.**
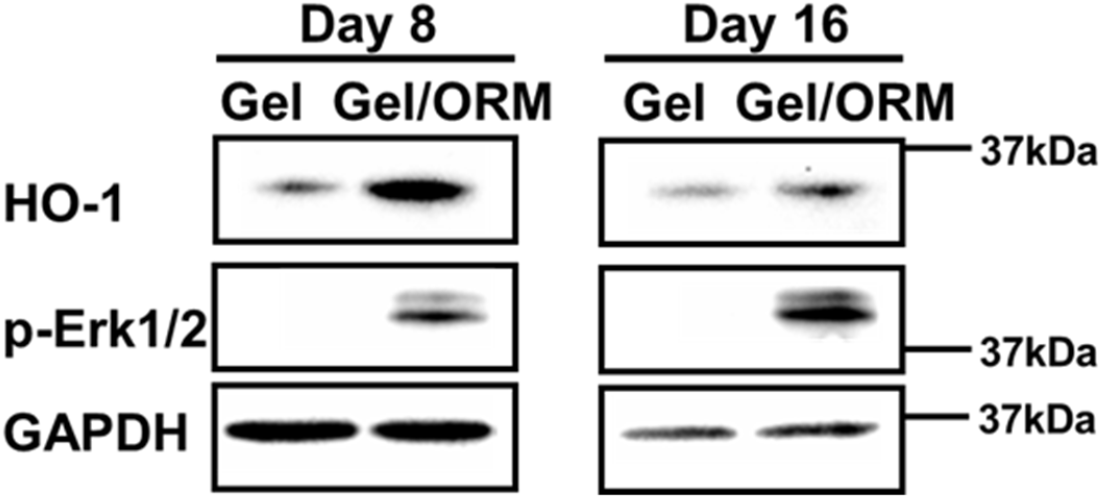
Oxygen-release microspheres activate HO-1 and Erk 1/2 pathways and promote pro-angiogenic growth factors expression during wound healing. Immunoblotting of HO-1 and phosphorylated Erk1/2 from the tissue lysates extracted from wounded skin at day 8 and 16 post surgery. GAPDH was used as a loading control.

## Discussion

Chronic hypoxia is a main characteristic of diabetic wounds and a serious impediment to the healing process. Consequently, sustained oxygenation of skin cells to mitigate chronic hypoxia represents an approach to accelerate diabetic wound healing. Oxygen therapy is advantageous over drug therapy as it has less concerns on controlling toxicity (*49*). However, current oxygen therapy approaches are unable to sustainedly provide sufficient oxygen to metabolic-demanding skin cells to promote diabetic wound healing (*50-53*). In this work, we developed an oxygen-release system to continuously oxygenate the diabetic wounds. It is composed of oxygen-release microspheres (ORMs), and their carrier, an injectable, thermosensitive, fast gelation, and ROS-scavenging hydrogel. The ORMs had core-shell structure with PVP/H_2_O_2_ complex as the core and a bioeliminable polymer as shell. The high molecular weight PVP/H_2_O_2_ complex reduces H_2_O_2_ diffusivity, allowing for sustained release of H_2_O_2_ during the hydrolysis of shell polymer (*54, 55*). The ORM surface was conjugated with catalase for timely converting H_2_O_2_ in the released PVP/H_2_O_2_ into molecular oxygen (**Fig. 1A**). Therefore, the ORMs were capable of directly releasing oxygen (**Fig. 1 E**). This is advantageous over current oxygen-release systems where most directly release H_2_O_2_ instead of oxygen (*24, 56, 57*). We demonstrated that the microspheres can release oxygen for at least 2 weeks (**Fig. 1E**), longer than most oxygen release systems (*24, 26*). The injectable, thermosensitive and fast gelation hydrogel was used as ORM carrier for largely retaining ORMs in the diabetic wounds upon delivery. The ROS-scavenging capability of the hydrogel serves two purposes, i.e., eliminating the H_2_O_2_ in the released PVP/H_2_O_2_ even if it is not completely converted by catalase, and capturing the upregulated H_2_O_2_ in diabetic wounds to decrease oxidative stress for accelerated wound healing. To the best of knowledge, current oxygen release system has not been designed to simultaneously release molecular oxygen and scavenge ROS.

We evaluated the efficacy and mechanism of action of the oxygen release system in the healing of excisional wounds in db/db mice (**Fig. 4A**). This model exhibits significant delay in wound closure and impaired wound bed vascularization compared with other well-accepted murine diabetes models, such as streptozocin-induced C57BL/6J and Akita (*58*). Our results show that the oxygen release system significantly promoted wound closure (**Fig. 4C**). We demonstrate that the accelerated wound healing is attributed to the sustained oxygenation and ROS scavenging in diabetic wounds that augmented cell survival, faster cell migration, stimulated angiogenesis, and reduced oxidative stress and inflammation (**Fig. 8**).

**Fig. 8.**
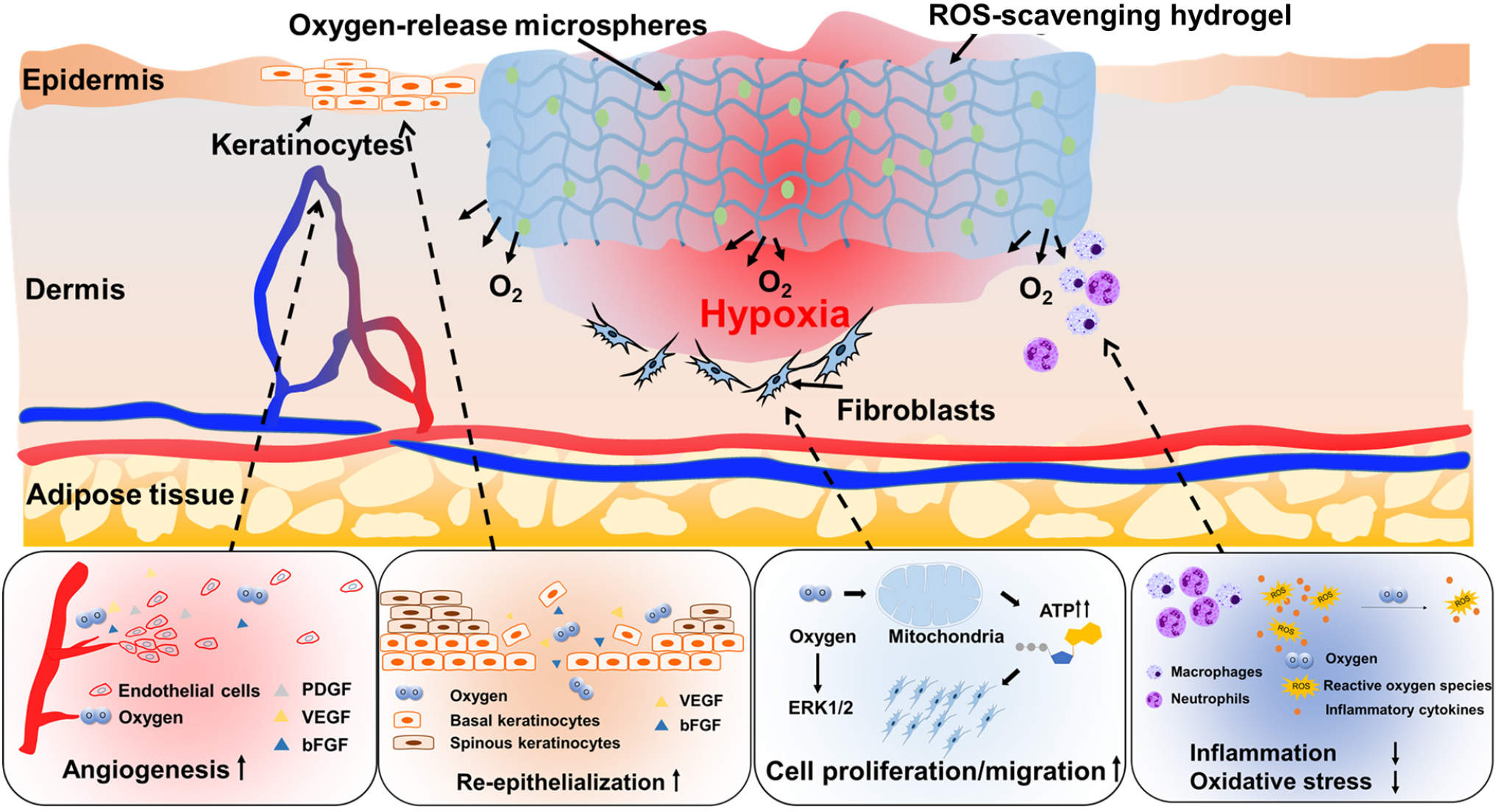
Mechanisms of accelerated wound healing by oxygen-releasing microspheres encapsulated in reactive oxygen species-scavenging hydrogel.

First, we investigated the effect of oxygen released from ORMs on skin cell survival under a hypoxic environment. We conducted in vitro studies under 1% oxygen and high glucose conditions to mimic the in vivo microenvironment in diabetic wounds (*11, 59*). The survival of keratinocytes, fibroblasts, and endothelial cells under hypoxia were significantly increased with the supplement of oxygen released from the ORMs (**Fig. 1, F to H**). These three cell types are responsible for epithelialization, wound contraction, and angiogenesis. In diabetic wounds, the treatment with oxygen-release system significantly increased the density of proliferating cells (**Fig. 5A, C**). The augmented cell survival can be attributed to the elevated cellular oxygen content (**Fig. 2C**). Cellular oxygen is essential for mitochondrial metabolism. The low oxygen level in the diabetic wounds leads to impaired mitochondrial metabolism (*60*). The release of oxygen significantly increased mitochondrial metabolism of the cells in the wounds evidenced by greater PGC1α+ cell density (**Fig. 5, B and D**). Mechanistically, we found that the enhanced skin cell survival by released oxygen is associated with the upregulation of p-Erk1/2 in vitro and in vivo (**Figs. 2E and 7**).

Next, we demonstrate that continuous oxygenation of skin cells promoted the migration and paracrine effects of keratinocytes and fibroblasts under hypoxia (**Fig. 1, L to Q**). The migration of these cells is essential for regeneration of epidermis and dermis. The enhanced paracrine effects in terms of upregulation of growth factors like *VEGFA, FGF2* and *PDGFB* may facilitate the regeneration. Specifically, *VEGFA* and *FGF2* have been found to increase keratinocyte migration and proliferation (*61-64*). The enhanced keratinocyte survival, migration and paracrine effects by continuous oxygenation led to significant increase of wound closure rate (**Fig. 4C**), and faster formation of the stratified epithelium (**Fig. 4D**).

Angiogenesis in diabetic wounds is essential for the regeneration of dermis. Although acute hypoxia induces angiogenesis mediated by hypoxia-inducible transcription factors, chronic oxygen deprivation cannot sustain this process, thereby impairing the healing process (*9*). We found that the released oxygen was able to increase endothelial cell survival (**Fig. 1H**) and tube formation (**Fig. 1, R to T**) in vitro. In the diabetic wounds, the continuous oxygenation significantly stimulated capillary formation (**Fig. 5, G and H**). The quicker angiogenesis may also be attributed to the enhanced paracrine effects as the expressions of angiogenic growth factors *Pdgfb* and *Vegfa* were substantially upregulated (**Fig. 5, E and F**).

Furthermore, we show that the developed oxygen delivery system alleviated oxidative stress in diabetic wounds (**Fig. 6, A and C**). The decreased oxidative stress was due to ROS-scavenging hydrogel since the ROS+ cell density in the Gel and Gel/ORM groups was similar. This eliminates potential concern that oxygen release may lead to ROS overproduction. The decreased oxidative stress may also be associated with increased HO-1 expression under hypoxic conditions (**Figs. 2E and 7**). HO-1 has antioxidant effects. It has been reported to promote diabetic wound healing (*65, 66*).

Lastly, we reveal that the oxygen-release system reduced inflammation in diabetic wounds (**Fig. 6, B and D**). Chronic wounds are characterized by a high concentration of pro-inflammatory M1 macrophages, the major contributor of pro-inflammatory cytokines, such as TNF-α, IL-1β, and CXCL1 (*67*). Additionally, hyperglycemia exacerbates inflammatory stress via production of pro-inflammatory cytokines like TNF-α (*68*). Our results show that the continuous oxygenation and ROS-scavenging greatly decreased the number of M1 macrophages and major pro-inflammatory cytokine expression (**Fig. 6, B, D, E and F**).

Overall, we demonstrate in this report that an oxygen-release system capable of simultaneously releasing oxygen and scavenging ROS significantly accelerated diabetic wound closure. The sustainedly released oxygen was multifunctional, i.e., promotion of skin cell survival, migration and paracrine effects, stimulation of endothelial tube formation and angiogenesis, and decrease of tissue inflammation. Compared with previous studies using growth factors or cells for the same animal model and wound assay, the oxygen-release system exhibited faster (*69-71*) or comparable wound closure rate (*72, 73*). Besides wound healing, the developed oxygen-release system may be used to treat other ischemic diseases such as peripheral artery disease, and coronary heart disease.

Our study has limitations. First, our study thus far is limited to rodent studies that may not be as representative of more complex human pathophysiology. Future studies will aim to apply our study tools in large animals. Second, the oxygen release kinetics and ROS-scavenging capability in diabetic wounds need to be optimized for different animal models. Third, to elucidate the underlying mechanisms that sustainedly released oxygen decreased inflammation in diabetic wounds, future mechanistic studies need to be conducted to reveal the associated signaling pathways. Nevertheless, the current study presents a new therapeutic approach for accelerated healing of chronic diabetic wounds without using drugs.

## Materials and Methods

### Materials

All chemicals were purchased from Millipore Sigma unless otherwise stated. NIPAAm (TCI) was recrystallized in hexane before use. 2-Hydroxyethyl methacrylate (HEMA, Alfa Aesar) was used before passing through a column filled with inhibitor removers. Benzoyl peroxide (BPO, Thermo Fisher Scientific), acryloyl chloride, 4-(hydroxymethyl)-phenylboronic acid pinacol ester, triethylamine (TEA, Fisher Scientific), PVP (40 kDa, Fisher Scientific), hydrogen peroxide solution (30%), bovine liver catalase (2000-5000 units/mg) were used as received.

### Synthesis and characterization of ROS-scavenging hydrogel

4-(Acryloxymethyl)-phenylboronic acid pinacol ester was synthesized by reaction between acryloyl chloride and 4-(hydroxymethyl)-phenylboronic acid pinacol ester (*74*). The ROS-scavenging hydrogel was synthesized by free radical polymerization of NIPAAm, HEMA, and 4-(acryloxymethyl)-phenylboronic acid pinacol ester using BPO as initiator, and 1,4-dioxane as solvent. The reaction was conducted at 70°C overnight with the protection of nitrogen. The polymer solution was precipitated in hexane. The polymer was purified twice by dissolving in tetrahydrofuran and precipitating in ethyl ether. Chemical structure of the polymer was confirmed by ^1^H-NMR (**Fig. S5**). The polymer was dissolved in Dulbecco’s phosphate-buffered saline (DPBS) at 4°C to make a 6% solution. The injectability of the 4 °C solution with or without 40 mg/mL oxygen-release microspheres was tested by a 26-gauge needle (*75, 76*).

Hydrogel H_2_O_2_ responsiveness was characterized by its weight loss after being incubated in H_2_O_2_ solution for 4 weeks. In brief, the hydrogel solution was solidified in 1.5 mL microcentrifuge tubes at 37 °C. After the supernatant was discarded, 0.2 mL, 37 °C DPBS with or without 50 mM H_2_O_2_ was added. The samples were collected at predetermined time points, and freeze-dried. The weight loss was calculated. To determine scavenging capability of the hydrogel for hydroxyl radicals and superoxide, Fenton reaction assay (*77*) and pyrogallol assay (*77, 78*) were performed, respectively. Briefly for Fenton reaction assay, hydrogel solution or DI water (control group) was incubated with FeSO_4_, Safranin O and H_2_O_2_ for 5 min, followed by heating at 55 °C for 30 min. After the samples were cooled to room temperature, the absorbance was measured at 492 nm using a microplate reader. For pyrogallol assay, the hydrogel solution or DI water was mixed with Tris-HCl. 3 mM pyrogallol solution was then added dropwise in the dark. The reaction was terminated by adding 8 M HCl. The absorbance was acquired at 299 nm.

### Cell survival on ROS-scavenging hydrogel in the presence of H_2_O_2_

The ROS-scavenging hydrogel was plated on 96-well plates. HaCaT cells (Addexbio) were seeded on the hydrogel surface at a density of 50000/well in optimized-DMEM (Addexbio) with 5% fetal bovine serum (FBS) and 1% penicillin-streptomycin. After 24 hours of culture, the medium was discarded. 200 μL of 100 μM H_2_O_2_-containing medium was added to each well. At 48 and 72 hours, the viability of HaCaT was measured by MTT assay (*79*). The non-ROS-responsive hydrogel, poly (NIPAAm-co-HEMA-co-acrylate-oligolactide), was used as control (*55, 76*).

### Fabrication of oxygen-release microspheres and catalase conjugation

The oxygen-release microspheres were fabricated by double emulsion method. Briefly, the shell of the microspheres was synthesized by the copolymerization of NIPAAm, HEMA, N-acryloxysuccinimide (NAS), and acrylate oligolactide (AOLA). The chemical structure of the shell polymer was confirmed by ^1^H-NMR (**Fig. S1**). The synthesized polymer was dissolved in dichloromethane to form the 5 wt% oil phase. The inner water phase was prepared by dissolving 242 mg of PVP in 1 mL, 30% H_2_O_2_ at 4°C overnight. The water phase was rapidly added into the oil phase, and sonicated by an ultrasonic liquid processor (Cole Parmer). The primary water-in-oil (W/O) emulsion was then poured into poly(vinyl alcohol) solution to form the water-in-oil-in-water (W/O/W) double emulsion. The mixture was stirred for 3 hours to remove dichloromethane, followed by centrifugation to collect the microspheres. Morphology and size of the microspheres were characterized by SEM images. To confirm the core-shell structure, fluorescein isothiocyanate (FITC) and rhodamine were added to the PVP/H_2_O_2_ solution and polymer/dichloromethane solution, respectively. The fluorescent images were acquired by a confocal microscope. To conjugate catalase onto the microsphere shell, 40 mg microspheres were mixed with 6 mL catalase solution (5 mg/mL in DI water), and stirred for 4 h at 4°C. The mixture was then centrifuged. The microspheres were washed 3 times with DI water to remove un-conjugated catalase. To confirm the conjugation, catalase was pre-labeled by FITC, and the fluorescent images of the microspheres were taken after the conjugation.

### Oxygen release kinetics

The ROS-sensitive hydrogel was dissolved in 4°C DPBS to make a 6 wt% solution. The oxygen-release microspheres were then mixed with the hydrogel solution at a concentration of 40 mg/mL. The oxygen release kinetics was determined by our established method (*54, 55, 76, 80*). Briefly, the wells of a 96-well plate were covered by a polydimethylsiloxane (PDMS) membrane loaded with an oxygen-sensitive luminophore Ru(Ph_2_phen_3_)Cl_2,_ and an oxygen-insensitive dye rhodamine B. 200 μL of mixture was then added into each well (n = 8 for each group). After gelation at 37°C, the supernatant was removed. 200 μL of DPBS pre-incubated in 1% oxygen condition was added. The oxygen release study was performed in 1% oxygen and 37°C conditions for 2 weeks. At each time point, the fluorescent intensity for Ru(Ph_2_phen_3_)Cl_2_ was measured (emission at 610 nm and excitation at 470 nm). The fluorescent intensity for rhodamine was also determined (emission at 576 nm and excitation at 543 nm), and used for normalization. Oxygen concentration was determined by the calibration curve.

### Cell survival, migration and ROS content under hypoxia

HaCaT cells (Addexbio) were cultured in optimized-DMEM (Addexbio) with 5% FBS and 1% penicillin-streptomycin. HDF (Lonza) were cultured in FGM-2 bulletkit (Lonza).HAEC (Cell Systems) were cultured in EGM-2 bulletkit (Lonza). The medium was changed every other day.

For all in vitro studies, cells were cultured using high glucose (450 mg/dL) and serum-free medium under 1% oxygen and 37°C conditions. To determine cell survival, the cells were cultured in a 96-well plate using medium with or without oxygen-release microspheres (40 mg/mL, n ≥ 5 for each group). The dsDNA content was measured using Picogreen dsDNA assay kit (Invitrogen) after 5 days of culture for HaCaT cells and HDFs, and after 7 days of culture for HAEC.

To perform cell migration assay, the cells were first cultured in a 6-well plate to reach 85-95% confluency (n = 4 for each group). The monolayer was then scraped with a 200 µL pipette tip, washed, and supplemented with serum-free medium with or without 40 mg/mL oxygen-release microspheres. After 48 hours, optical images were taken by an optical microscope (Olympus IX70). The distances between two sides of the scratch were measured by ImageJ. The migration ratio was calculated as:

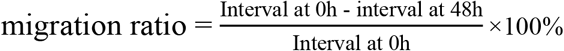

To measure intracellular ROS content, the cells were pre-stained with ROS sensitive dye CM-H_2_DCFDA, and cultured with serum-free medium with or without 40 mg/mL oxygen-release microspheres. After 3 days of culture for HaCaT cells and HAEC, and 5 days of culture for HDF, the cells were fixed and stained with DAPI. Fluorescent images were taken using a confocal microscope. The CM-H_2_DCFDA positive cell density was quantified from at least 10 images for each group, and then normalized to the CM-H_2_DCFDA positive cell density cultured under normoxia.

### Measurement of intracellular oxygen content

To determine the intracellular oxygen content, HaCaT cells were incubated with lithium phthalocyanine (LiPc) nanoparticles for 2 hours to allow cellular uptake. The residual nanoparticles were washed with DPBS for 3 times. After trypsinization, the cells were encapsulated in the ROS-sensitive hydrogel or mixture of hydrogel and oxygen-release microspheres (40 mg/mL). The samples were transferred into EPR tubes (Wilmad-LabGlass, n = 3 for each group). The EPR tubes were opened on both sides and placed in a hypoxic incubator (1% oxygen, 37°C) for 4 hours for complete gelation and gas balance. After that, the tubes were sealed and incubated for 24 hours under 1% oxygen. The EPR spectrum was recorded using an X-band EPR instrument (Bruker). The parameters used in this experiment were 0.1 mW for microwave power, 1.0 db for attenuation and 9.8 GHz for frequency, following our reported method (*80, 81*). Oxygen partial pressure (pO_2_) was calculated from the linewidth of the spectrum, and calibration curve of linewidth vs. oxygen concentration.

### Measurement of intracellular ATP level

To determine the intracellular ATP level, HaCaT were cultured in a 6-well plate to reach 85-95% confluency. The culture media was then replaced with serum-free media with or without oxygen-release microspheres (40 mg/mL). After 24-hour incubation under 1% oxygen, the cells were lysed and the ATP content was measured by an ATP assay kit per manufacturer’s instruction (Millipore Sigma, n = 3).

### Endothelial tube formation

HAECs were cultured in a 3D collagen model described previously (*28*). Briefly, the model was prepared by mixing rat tail collagen type I (Corning), FBS, DMEM and NaOH. 400 uL of the mixture was placed in a 48-well plate and incubated at 37°C for 30 min to allow the formation of jelly-like solid collagen gel. HAECs were encapsulated in ROSS gel solution and injected into the collagen-based 3D model at a density of 10000 cells/well. After a 24-hour hypoxic culture, HAECs were fixed by 4% paraformaldehyde, and stained with DAPI and F-actin. Fluorescent images were taken as z-stacks with a confocal microscope (Zeiss LSM700).

### Implantation of hydrogel and oxygen-release microspheres into diabetic wounds

All animal care and experiment procedures were conducted in accordance with the National Institutes of Health guidelines. The animal protocol was approved by the Institutional Animal Care and Use Committee of Washington University in St. Louis. 8-week old, female db/db mice (BKS.Cg-Dock7m +/+ Leprdb/J) were purchased from Jackson Laboratory. Blood glucose was measured before surgery to ensure that the glucose level was greater than 300 mg/dL. After anesthetization and removal of hair, 2 full-thickness, 5 mm-diameter wounds on the dorsal skin were created for each mouse using a biopsy punch. The hydrogel (6% in DPBS) or Gel/ORM (6% in DPBS, 40 mg/mL microspheres) was topically administrated. Digital images of the wounds (n≥ 8 for each group) were taken every other day, and the wound size was determined by ImageJ. After 8 and 16 days of implantation, the mice were sacrificed. The wound tissues were collected for histological and immunohistochemical analysis, and RNA and protein isolation.

### Histological and immunohistochemical analysis

The wound tissues were fixed in 4% paraformaldehyde overnight, embedded in paraffin, and serially sectioned at 5 µm. H&E staining was performed to measure the epidermal thickness in wounded skin. Picrosirius red staining was used to quantify the collagen deposition and collagen I/III ratio. For immunohistochemical analysis, following deparaffinization, antigen retrieval and blocking, the sections were incubated with primary antibodies including rabbit monoclonal anti-cytokeratin 10 (K10, Abcam), mouse monoclonal anti-cytokeratin 14 (K14, Abcam), rabbit monoclonal anti-CD86 (Abcam), rat monoclonal anti-Ki67 (Invitrogen), isolectin GS-IB_4_ (Invitrogen) (*82, 83*), and ROS indicator (CM-H_2_DCFDA, Invitrogen). The sections were then incubated with corresponding secondary antibodies. Nuclei were stained with DAPI. Quantification of the stainings was performed using ImageJ. At least 6 images for each group were used for the quantification.

### mRNA expression in the in vitro cultured cells and diabetic wounds

RNA was isolated from the in vitro cultured cells or wound tissues by TRIzol following manufacturer’s instruction. cDNA was synthesized using high-capacity cDNA reverse transcription kit (Thermo Fisher). Gene expression was performed by real-time RT-PCR using SYBR green (Invitrogen) and selected primer pairs (**Table S1**). β-actin was served as a housekeeping gene. Data analysis was performed using the ΔΔCt method.

### Protein expression in the in vitro cultured cells and diabetic wounds

Protein lysates were collected from the in vitro cultured cells or wound tissues, and separated by SDS-PAGE. LF-PVDF membranes (Bio-rad) were used to transfer the proteins at 4 °C overnight, followed by blocking and incubation with primary antibodies at 4 °C. The blots were washed with PBST (DPBS+0.1% Tween 20) and incubated with appropriate HRP-conjugated secondary antibodies. Immunoblots were detected by a detection kit from Advansta and imaged using ChemiDoc XRS+ System (Bio-rad). The primary antibodies used were anti-GAPDH (1:4000, Abcam), anti-HO-1 (1:500, Abcam), anti-phospho-Erk1/2 (1:500, Cell signaling), and anti-Erk1/2 (1:1000, Cell signaling).

For protein array assay, wounded tissue lysates were collected, and tested using Proteome profiler mouse cytokine array kit (R&D Systems) according to the manufacturer’s instructions. The pixel density was quantified by Image Lab software (Biorad) (*84*).

### Statistical analysis

All data were presented as mean ± standard deviation. Statistical analysis was performed between groups using one-way ANOVA with Tukey’s post hoc test. A value of p < 0.05 was considered statistically significant.

## Acknowledgements

Confocal images were performed in part through the use of Washington University Center for Cellular Imaging (WUCCI) supported by Washington University School of Medicine, The Children’s Discovery Institute of Washington University and St. Louis Children’s Hospital (CDI-CORE-2015-505 and CDI-CORE-2019-813) and the Foundation for Barnes-Jewish Hospital (3770 and 4642). We thank Musculoskeletal Research Center in Washington University School of Medicine for the help with histological sections. We also thank Dr. Manmilan Singh from Department of Chemistry in Washington University for his help with EPR test.

## Author contributions

J.G., Y.G. and H.N. designed the study. Y.G., H.N., and Z.L. performed most of the experiments. J.G., Y.G., H.N., and Y.D. analyzed the results and wrote the manuscript. M.Z., L.M., and J.S. provided insights of the study and edited the manuscript. J.G. supervised the project.

## Data and materials availability

All data needed to evaluate the conclusions in the paper are present in the paper and/or the Supplementary Materials. The data that support the study may be requested from the corresponding author.

