## Supplementary material for "Sustained Oxygenation Accelerates Diabetic Wound Healing by Simultaneously Promoting Epithelialization and Angiogenesis, and Decreasing Tissue Inflammation": Figs. S1 to S5, Table S1

**This file includes:**

Figs. S1 to S5

Table. S1

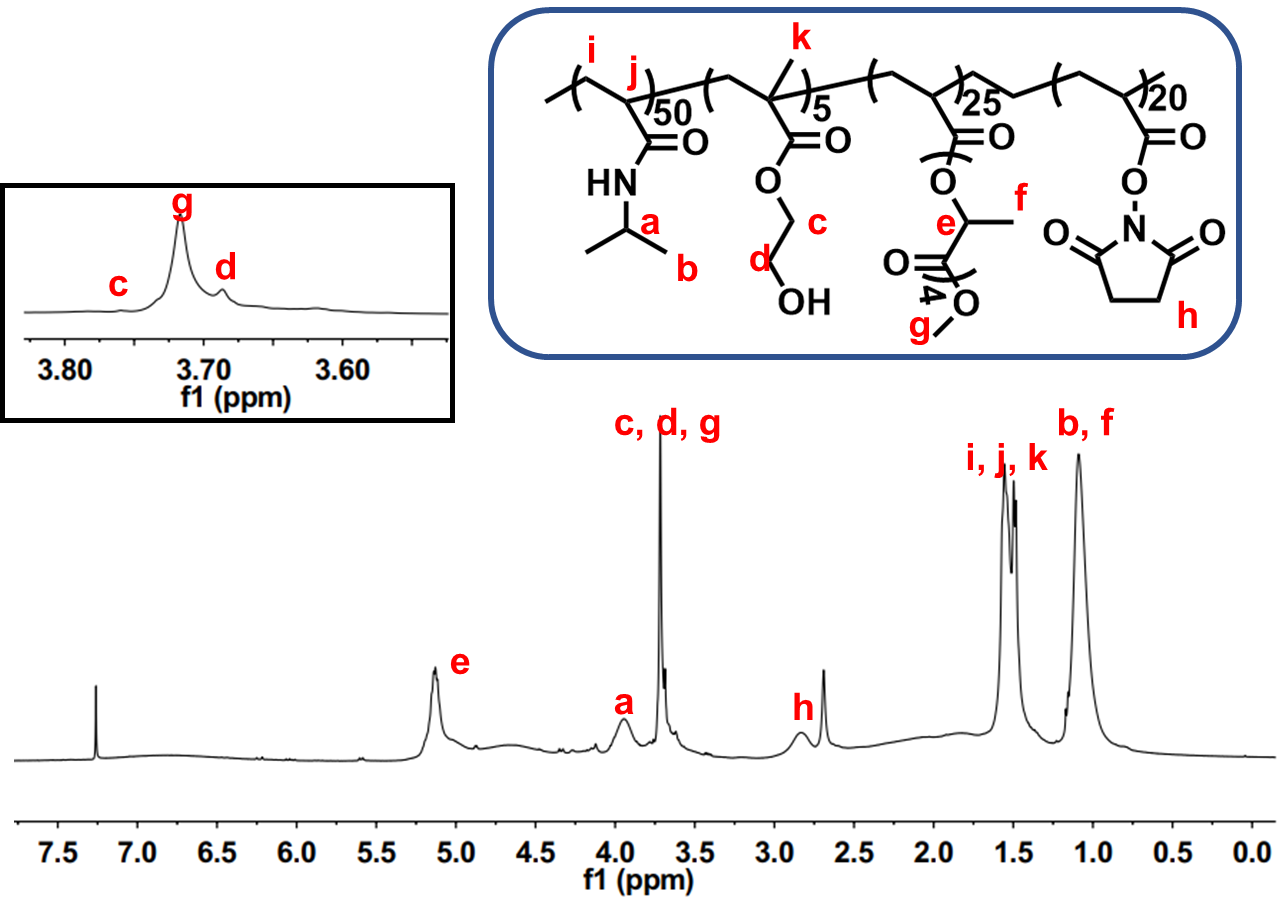
Fig. S1.

^1^H-NMR of poly (NIPAAm-co-HEMA-co-AOLA-co-NAS)

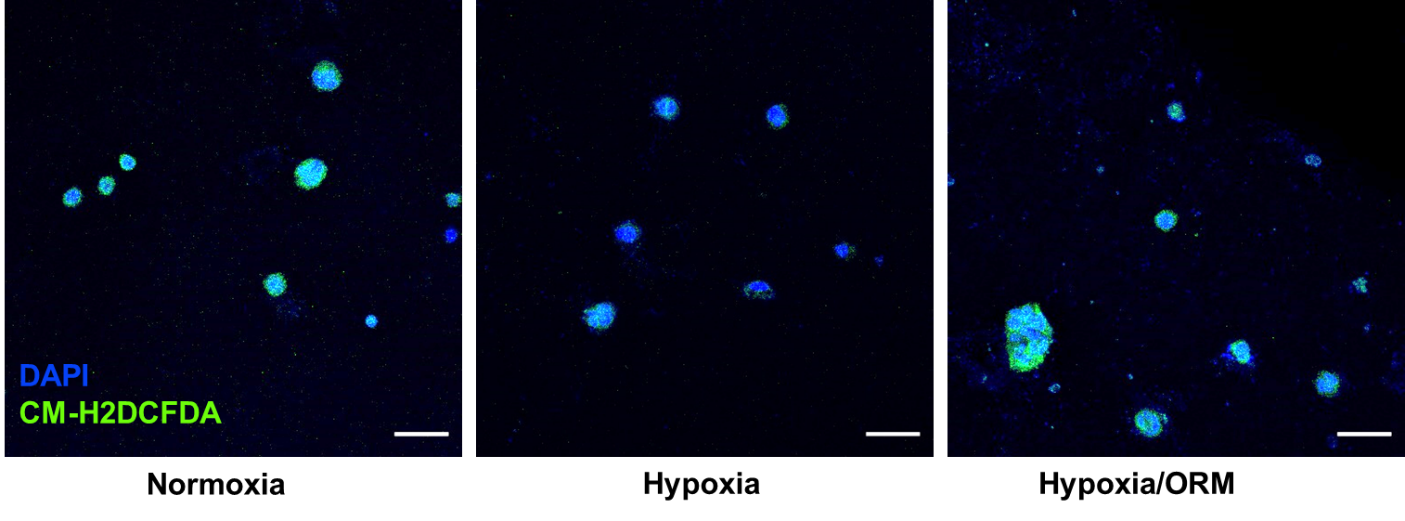

Fig. S2.

Fluorescent images of HaCaT cells stained with CM-H_2_DCFDA (green) and DAPI cultured under normoxia or hypoxia for 3 days.

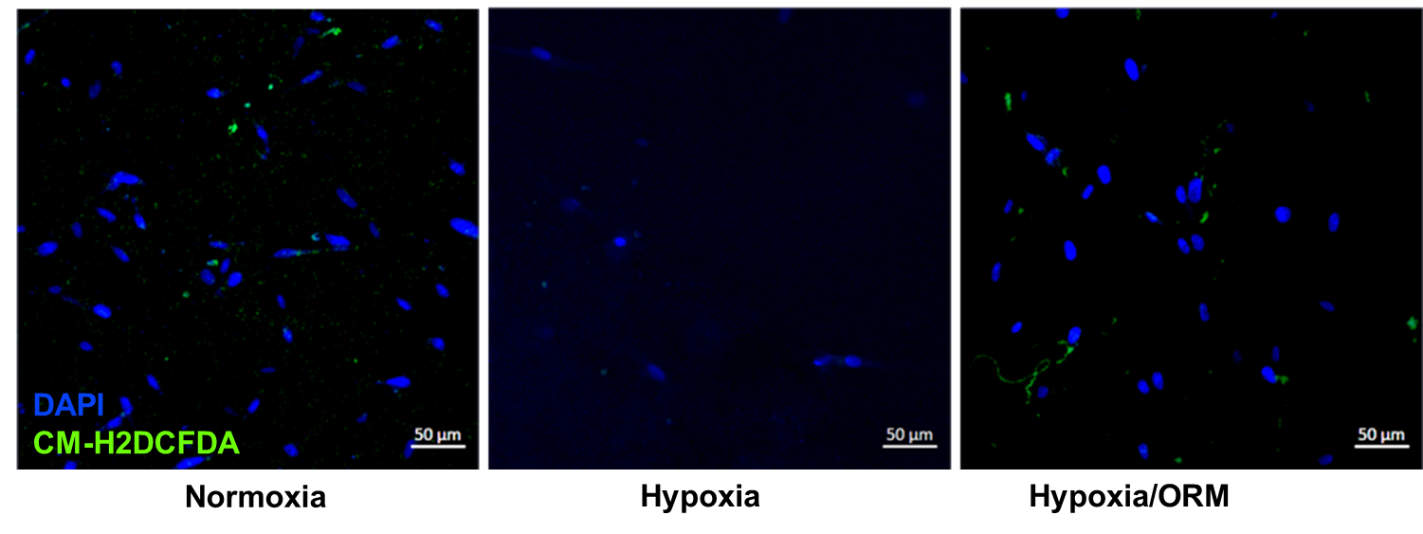

Fig. S3.

Fluorescent images of HDF stained with CM-H2DCFDA (green) and DAPI cultured under normoxia or hypoxia for 5 days.

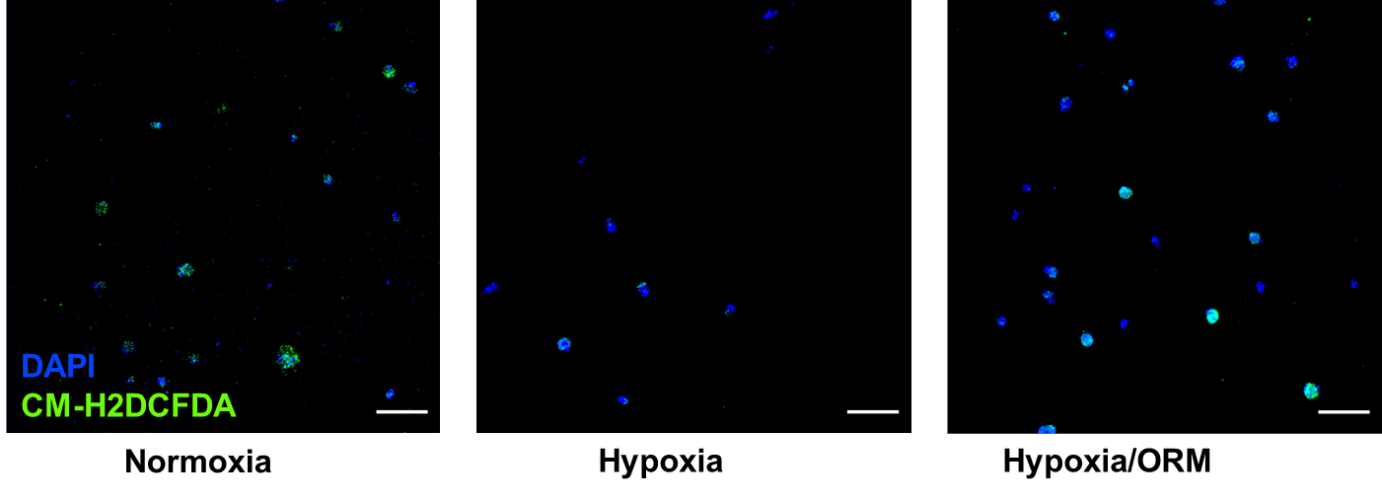

**Fig. S4.**

Fluorescent images of HAEC stained with CM-H_2_DCFDA (green) and DAPI cultured under normoxia or hypoxia for 3 days

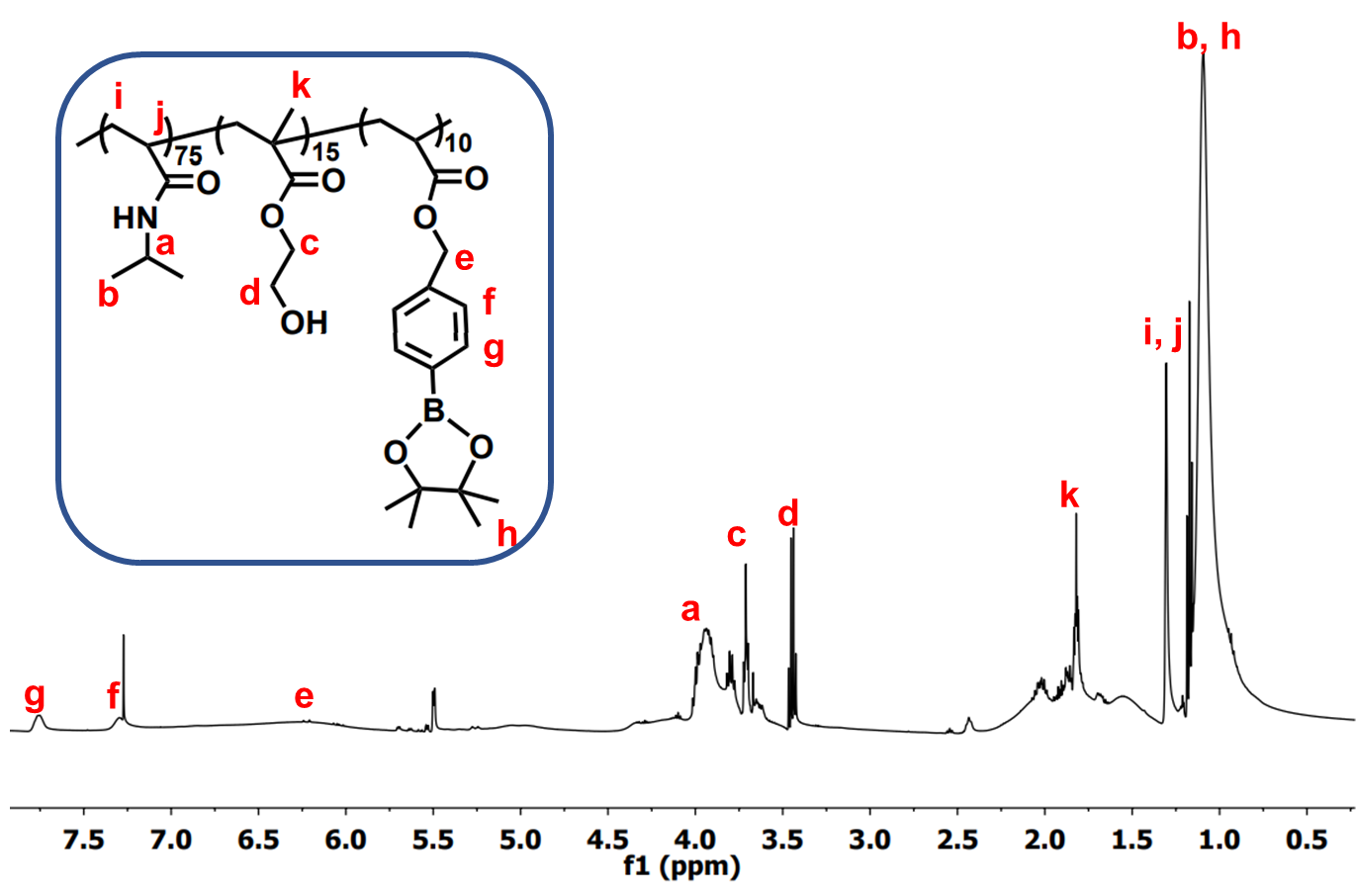

**Fig. S5.**

^1^H-NMR of poly(NIPAAm-co-HEMA-co-4-(acryloxymethyl)-phenylboronic-acid-pinacol-ester)

Table S1.

List of primer sequences used in real-time RT-PCR for in vitro and in vivo studies.

| Gene | Forward (5’-3’) | Reverse (5’-3’) | Species |
| --- | --- | --- | --- |
| *PDGFB* | GGGCAGGGTTATTTAATATGG | AATCAGGCATCGAGACAG | Human |
| *VEGFA* | AATGTGAATGCAGACCAAAG | GACTTATACCGGGATTTCTTG | Human |
| *FGF2* | TGGCTTCTAAATGTGTTACG | GTTTATACTGCCCAGTTCG | Human |
| *Pdgfb* | GTGGGCAGGGTTATTTAATATG | GAGGGGAACAACATTATCAC | Mouse |
| *Vegfa* | TAGAGTACATCTTCAAGCCG | TCTTTCTTTGGTCTGCATTC | Mouse |
